# Escape from mitotic catastrophe by actin-dependent nuclear displacement in fission yeast

**DOI:** 10.1101/2020.12.03.411074

**Authors:** Masashi Yukawa, Yasuhiro Teratani, Takashi Toda

## Abstract

Proper nuclear positioning is essential for the execution of a wide variety of cellular processes in eukaryotic cells (Gundersen and Worman, 2013; Kopf et al., 2020; Lele et al., 2018). In proliferating mitotic cells, nuclear positioning is crucial for successful cell division. The bipolar spindle, which pulls sister chromatids towards two opposite poles, needs to assemble in the geometrical center of the cell. This ensures symmetrical positioning of the two nuclei that are reformed upon mitotic exit, by which two daughter cells inherit the identical set of the chromosomes upon cytokinesis. In fission yeast, the nucleus is positioned in the cell center during interphase; cytoplasmic microtubules interact with both the nucleus and the cell tips, thereby retaining the nucleus in the medial position of the cell (Daga et al., 2006; Tran et al., 2001). By contrast, how the nucleus is positioned during mitosis remains elusive. Here we show that several cell-cycle mutants that arrest in mitosis all displace the nucleus towards one end of the cell axis. Intriguingly, the actin cytoskeleton, not the microtubule counterpart, is responsible for the asymmetric movement of the nucleus. Time-lapse live imaging indicates that mitosis-specific F-actin cables interact with the nuclear membrane, thereby possibly generating an asymmetrical pushing force. In addition, constriction of the actomyosin ring further promotes nuclear displacement. This nuclear movement is beneficial, because if the nuclei were retained in the cell center, subsequent cell division would impose the lethal cut phenotype (Hirano et al., 1986; Yanagida, 1998), in which chromosomes are intersected by the contractile actin ring and the septum. Thus, fission yeast escapes from mitotic catastrophe by means of actin-dependent nuclear movement.

## RESULTS AND DISCUSSION

### Mitotic arrest leads to nuclear displacement

Cut7 in fission yeast belongs to the Kinesin-5 family and plays an essential role in bipolar spindle assembly (Hagan and Yanagida, 1990; Yukawa et al., 2020). Its inactivation leads to the emergence of monopolar spindles and mitotic arrest. Revisiting defective phenotypes of *cut7* temperature-sensitive (ts) mutant cells indicated that chromosomes were often displaced from the cell center upon incubation at 36°C, by which septated cells contained one compartment with chromosomes, while the other compartment was anucleate (Figure 1A). On the other hand, cells whose nuclei were retained in the middle displayed the “cut” phenotype (Hirano et al., 1986; Yanagida, 1998), in which chromosomes are intersected by the septum.

**Figure 1.**
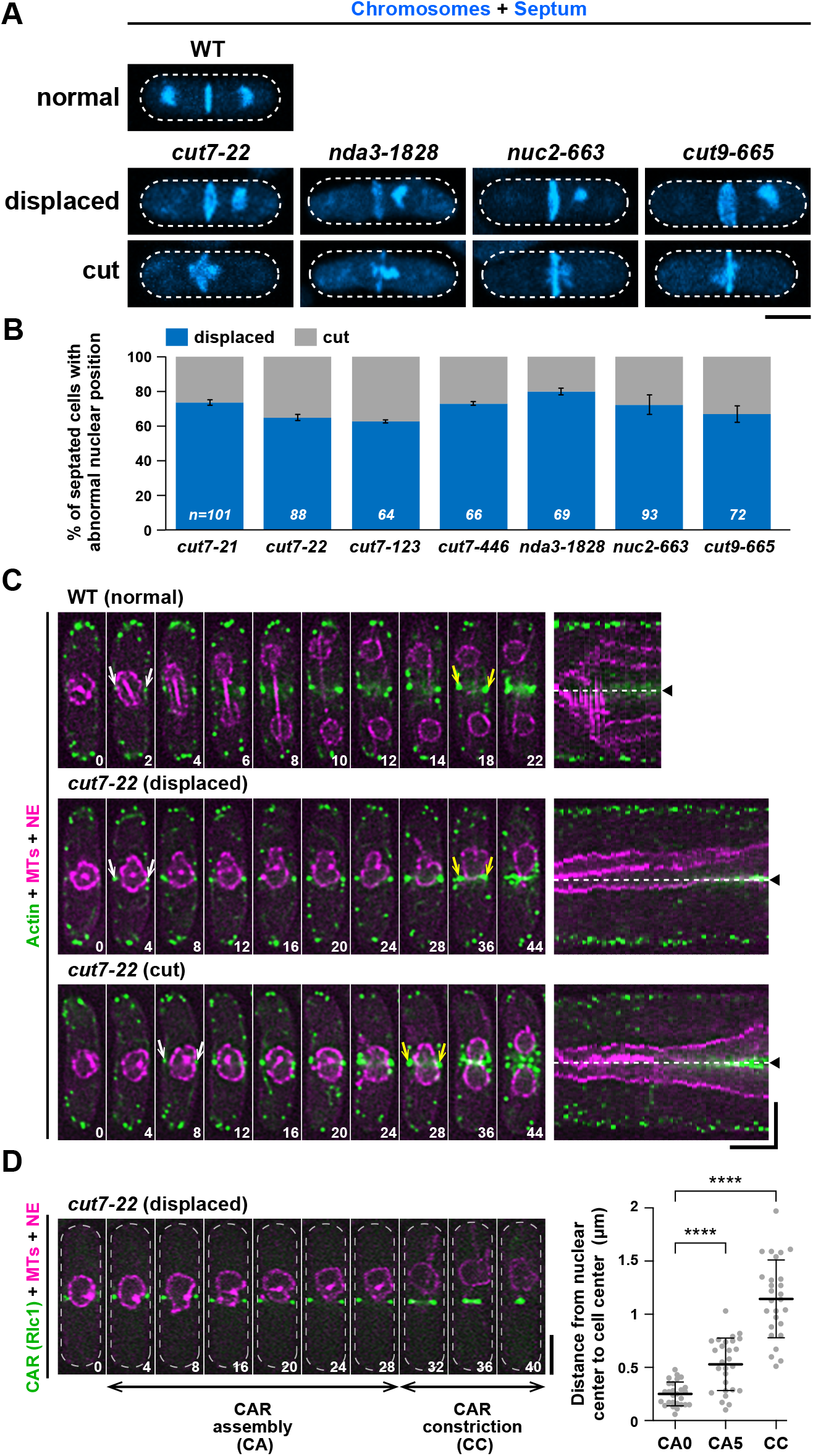
Mitotic mutants exhibit nuclear displacement. (A) Nuclear displacement in several mitotic mutant cells. Exponentially growing wild type or indicated mitotic mutants grown at 27°C were shifted to 36°C and incubated for 3 h except for *nda3-1828*, which was incubated for 6 h. Cells were fixed with methanol, and stained with DAPI (chromosomes) and Calcofluor (septa). Representative cells for wild type (top row), mutants displaying displaced chromosomes (middle row) or cut (bottom row) are shown. Scale bar, 5 μm. (B) The percentage of cells showing displaced chromosomes or cut. The sample numbers (n) for individual strains are indicated on the bottom of columns. Data are presented as the means ± S.D. (C) Time-lapse and kymograph images. Wild type (top row) or *cut7-22* cells (middle and bottom rows) grown at 27°C were shifted to 36°C and incubated for 2 h, when time-lapse imaging started. Cells contain mCherry-Atb2 (magenta, MTs), Cut11-mRFP (magenta, the NE) and LifeAct-GFP (green, actin). The first time-points when LifeAct-GFP signals were observed in the middle of cells are indicated with white arrows, while those when the CAR initiated constriction are marked with yellow arrows. Corresponding kymographs are shown on the right (one minute interval images were merged), in which the middle of the cell axis is shown with dotted lines and arrowheads. Scale bars, 10 min (horizontal) and 5 μm (vertical). (D) Timing of nuclear displacement in relation to that of CAR assembly and constriction. *cut7-22* cells containing mCherry-Atb2 (magenta, MTs), Cut11-mRFP (magenta, the NE) and Rlc1-GFP (green, the CAR) were grown at 27°C and shifted to 36°C for 2 h, when time-lapse imaging started (left). The duration of CAR assembly (CA) and constriction (CC) are marked with horizontal arrow bars. Scale bars, 5 μm. On the right graph, the position of the nucleus (distance from the nuclear center to cell center) is plotted against initiation of CAR assembly (CA0), 5 min after CAR assembly (CA5) and initiation of CAR constriction (CC). All *p*-values were obtained from the two-tailed unpaired Student’s t test. **** *p* <0.0001. The numbers on the bottom right corner of each image show times in minutes (C and D).

Nuclear displacement is not specific to Kinesin-5 malfunction, as we observed the same phenotypes in other mitotic mutants including mutations in β-tubulin (*nda3-1828*) (Radcliffe et al., 1998) and two subunits of the Anaphase Promoting Complex/Cyclosome (APC/C) (*nuc2-663* and *cut9-665*) (Hirano et al., 1988; Samejima and Yanagida, 1994; Yamada et al., 1997) (Figure 1A and 1B). It is of note that the displacement of the nucleus in *nuc2-663* and *cut9-665* mutants was previously noted (Yanagida, 1998). As fungi undergo a closed mitosis, the displacement of the chromosomes would stem from a defect in nuclear positioning. As a representative, we used *cut7-22* for the following investigation, unless otherwise stated.

### The nucleus becomes off-center as the medial actomyosin ring assembles

We examined the dynamics of nuclear movement using time-lapse fluorescence microscopy. For this purpose, wild type and *cut7-22* strains were tagged with fluorescent markers for the nuclear envelope (NE, Cut11-mRFP; Cut11 is also localized to the mitotic spindle pole body (SPB) (West et al., 1998)), microtubules (MTs, mCherry-Atb2; α2-tubulin (Toda et al., 1984)) and actin (LifeAct-GFP (Huang et al., 2012)) or Type II myosin regulatory light chain (Rlc1-GFP (Le Goff et al., 2000; Naqvi et al., 2000)). In fission yeast, the contractile actomyosin ring (CAR) initiates assembly during early mitosis, matures into a complete ring and then constricts in telophase, followed by cytokinesis (a schematic illustration is shown in Figure S1A and time-lapse images in Figure 1C; Video S1). Live imaging of *cut7* mutant cells incubated at the restrictive temperature indicated that there were two populations as described earlier (Figure 1B). The first type (30/57) exhibited nuclear displacement (Figure 1C; Video S2). The second type (27/57) represented ellipsoidal nuclei with the CAR being formed in the middle of the cell axis destined for the cut phenotype (Figure 1C; Video S3). Interestingly, the timing of nuclear displacement coincided with or was close to that of CAR assembly prior to CAR constriction (Figure 1D; Video S4).

We addressed whether mitotic delay is necessary for nuclear displacement. For this purpose, the *mad2* gene, encoding a core component of the spindle assembly checkpoint (SAC) (Musacchio, 2015), was deleted in *cut7-22*. The degree of nuclear displacement was lessened; however, we still observed nuclear movement in *cut7-22mad2Δ* cells (Figure S1B). Therefore, during prolonged mitotic arrest, the nucleus moves towards one end of the cell as these cells undergo CAR assembly and this could happen without mitotic delay.

### Nuclear displacement depends upon the F-actin cytoskeleton

We wished to identify the mechanism by which nuclear displacement is elicited. As the nucleus moves in accordance with CAR assembly, we treated *cut7* cells with an F-actin depolymerizing drug, Latrunculin A (LatA) at 36°C. Remarkably, LatA-treated *cut7* mutant cells almost completely ceased nuclear displacement (Figure 2A; Video S5). Figure 2B shows the kinetics of nuclear movement in the absence or presence of LatA. In all cases examined (n=12), LatA treatment displayed very minimal nuclear fluctuations (Figure 2C). By contrast, in the absence of LatA (n=18), ~44% or ~11% of cells displayed the maximal distance of >1 μm or 0.5-1.0 μm, respectively (Figure 2C). We also treated *nda3-1828*, *nuc2-663* and *cut9-665* cells with LatA at 36°C, and found that as in the case for *cut7*, nuclear displacement was suppressed in these cells by actin depolymerization (Figure S1C and S1D). These results indicated that the F-actin cytoskeleton is responsible for mitotic nuclear displacement and suggested that the driving force is generated by the polymerized actin structures.

**Figure 2.**
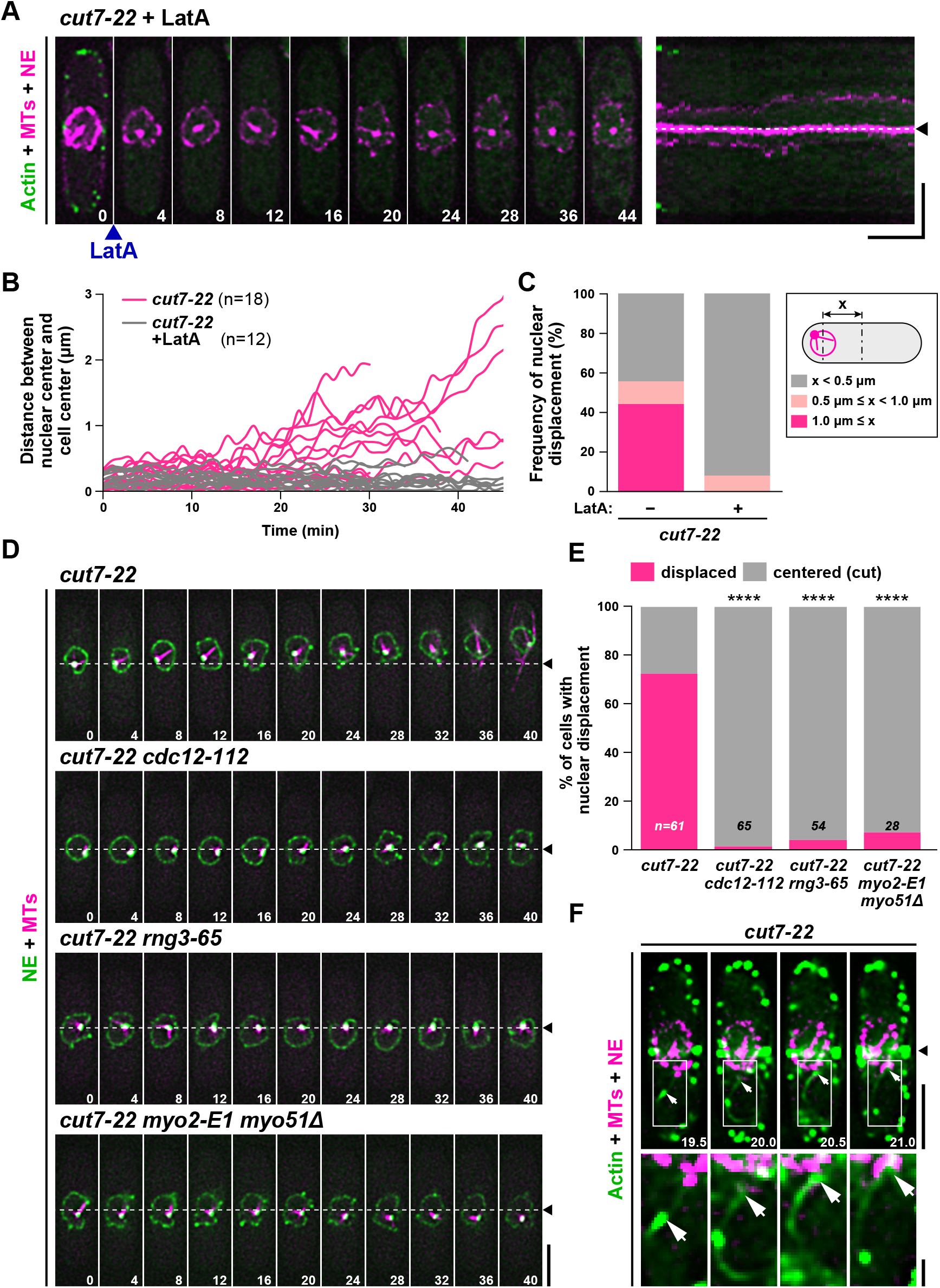
Cdc12/Formin, Myo2 and Myo51 are responsible for nuclear displacement. (A) Time-lapse and kymograph images of the nuclear position in *cut7-22* cells incubated at 36°C in the presence of LatA. *cut7-22* mutant cells grown at 27°C were shifted to 36°C and incubated for 2 h (time 0). At this point, LatA (50 μM) was added and time-lapse imaging started. Cells contained mCherry-Atb2 (magenta, MTs), Cut11-mRFP (magenta, the NE) and LifeAct-GFP (green, actin). Note that actin signals already disappeared 4 min after LatA treatment. Scale bars, 10 min (horizontal) and 5 μm (vertical). (B) Profiles of the relative position of the nucleus in *cut7-22* mutants in the absence (n=18) or presence (n=12) of LatA. The distance (x μm) between the center of the cell axis and that of the nucleus was measured at each time-point and plotted against time. (C) The degree of nuclear movement in *cut7-22* cells in the absence or presence of LatA. The percentage of cells that do or do not show nuclear displacement is shown. For each sample, the maximal distance (x μm) between the center of the cell axis and that of the nucleus was determined using the data shown in (B) and categorized into three classes: displaced (shown in magenta, in which x is ≥1 μm), mildly displaced (shown in pink, in which x is between 0.5 μm and 1 μm) and centered (shown in gray, in which x is ≤0.5 μm). (D) Time-lapse images of the nuclear position. Indicated mutant cells grown at 27°C were incubated at 36°C for 2 h, when live imaging started. Individual mutants contain mCherry-Atb2 (magenta, MTs) and Cut11-GFP (green, NE). Scale bar, 5 μm. (E) Profiles of the relative position of the nucleus in individual mutants. If x (as in (C)) was <1 μm or a cell showed cut, it was classified as centered (cut). If x was >1 μm, it was classified as displaced. The sample numbers are shown on the bottom of each column. All *p*-values were obtained from the two-tailed χ^2^ test. **** *p* <0.0001. (F) Visualization of interaction between F-actin cables and the NE in *cut7-22*. A *cut7-22* strain used in (A) was grown at 27°C and incubated at 36°C for 2 h, when time-lapse imaging started. The bottom row shows enlarged images in squares shown on the top row. Arrows point the tips of F-actin cables (green) that interact with the NE (magenta). Scale bars, 5 μm (top) and 1 μm (bottom). The position of the cell center is shown with dotted lines and arrowheads (A, D) or arrowhead (F) and the numbers on the bottom right indicate times in minutes upon recording (A, D and F).

### Cdc12/Formin, but not For3/Formin, is required for nuclear displacement

In fission yeast, F-actin assembles into three different structures during vegetative growth cycles: filamentous actin cables, endocytic actin patches and the CAR (Kovar et al., 2011). To distinguish which actin structures are responsible for force generation, we first treated *cut7-22* cells with CK-666, a small molecule that specifically inhibits the Arp2/3 complex, thereby disassembling actin patches (Burke et al., 2014; Nolen et al., 2009). Addition of this inhibitor resulted in the loss of actin patches as expected (Figure S2A). Nonetheless, nuclear displacement still occurred (Figure S2B and S2C). Notably, the degree of nuclear displacement was augmented by CK-666 treatment. This indicates that endocytic actin patches are not essential for force generation.

Next, we used the *cdc12-112* ts mutant; *cdc12* encodes Formin that is required for assembly of mitotic actin cables and the CAR (Chang et al., 1997; Huang et al., 2012). We constructed *cut7-22cdc12-112* double mutants, which were indeed defective in actin cable assembly and CAR formation (Figure S2D). In this background, nuclear movement was almost completely suppressed (Figure 2D and 2E; Video S6), indicating that the absence of nuclear displacement is a consequence of the lack of mitosis-specific actin cables, which are required to assemble the CAR. We then addressed the roles of myosin. Using the *rng3-65* ts mutant (Lord and Pollard, 2004; Wong et al., 2000), which is defective in myosin assembly and activation, we found that the nucleus remained in the middle of *cut7-22rng3-65* mutant cells (Figure 2D and 2E; Video S7). Furthermore, a combination of mutations in myosins II and V (*myo2-E1* and *myo51* deletion) supressed nuclear displacement in *cut7-22* (Figure 2D and 2E; Video S8). This result is totally in line with the previous data showing that Myo2 and Myo51 together corroborate to form proper mitotic actin cables and the CAR (Huang et al., 2012). Myo1 and Myo52 (another myosin V) appear unimportant, as *cut7-22myo1Δ* and *cut7-22myo2-E1myo52Δ* cells exhibited nuclear displacement (Figure S2D and S2E). In clear contrast to the requirement of Cdc12, another Formin For3, which promotes actin cable formation mainly during interphase (Feierbach and Chang, 2001), is dispensable for nuclear movement (Figure S2E and S2F).

Close inspection of time-lapse movies during short intervals captured time-points at which actin cables interacted with the NE in the *cut7-22* mutant (Figure 2F; Video S9). While it is possible that the nucleus is displaced poleward through myosin-mediated pulling force (Lo Presti et al., 2012), as interaction between actin cables and the nuclear membrane appears transient, we hypothesize that mitosis-specific F-actin cables push the nucleus, leading to nuclear displacement upon prolonged mitotic arrest.

### Actomyosin ring constriction accelerates nuclear displacement

Although the nucleus initiates displacement coincident with CAR assembly prior to its constriction (see Figure 1D), the process of ring closure might cooperate to promote nuclear movement. To scrutinise this possibility, we constructed double mutants between ts *cut7-22* and *cdc7-24* (Nurse et al., 1976). The Cdc7 kinase is a component of the SIN (Septation Initiation Network) that is required for not only septum formation but also maturation and constriction of the CAR (Krapp and Simanis, 2008; Roberts-Galbraith and Gould, 2008). Upon incubation of *cut7-22cdc7-24* mutant cells at 36°C, nuclear movement was imaged with time-lapse microscopy. As shown in Figure 3A and 3B, the degree of nuclear movement was substantially suppressed. Nonetheless, we noticed residual movement (Figure 3C; Video S10). Next, we added LatA to *cut7-22cdc7-24* mutant cells upon shift to 36°C. Interestingly, LatA almost completely halted nuclear movement (Figure 3A-3C; Video S11). Taken together, we posit that nuclear displacement is induced initially by a force generated through F-actin cables that incorporate into the CAR and then the movement is further accelerated by a force derived from CAR constriction.

**Figure 3.**
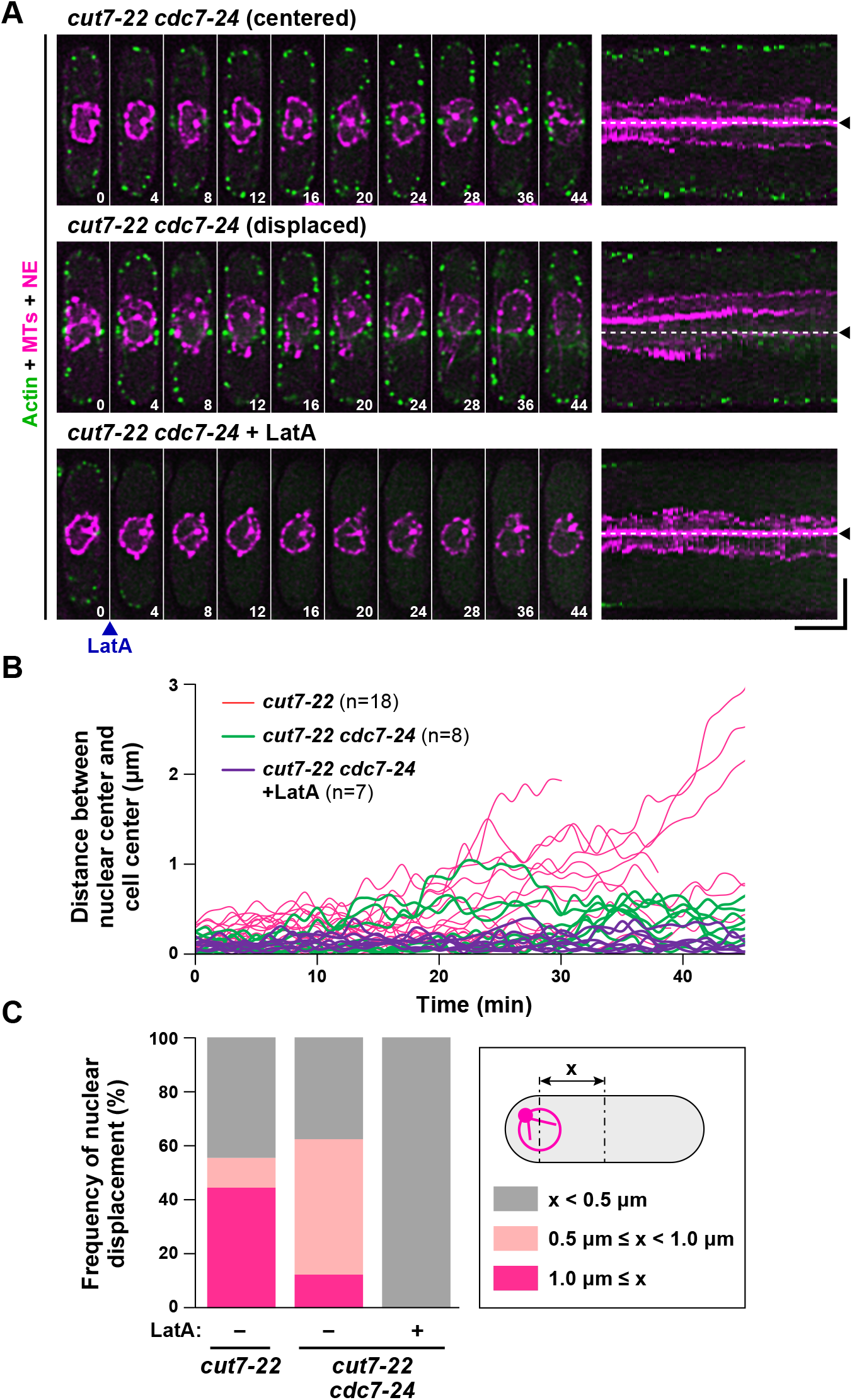
Inhibition of actomyosin ring constriction substantially but not completely abolishes nuclear displacement. (A) Time-lapse and kymograph images of the nuclear position in *cut7-22cdc7-24* cells incubated at 36°C in the absence or presence of LatA. Mutant cells were grown at 27°C, shifted to 36°C and incubated for 2 h, when imaging started. Two representative cells, in which the nucleus remains centered (top row) or is displaced (middle row), are shown. LatA (50 μM) was added at time 0. Cells contained mCherry-Atb2 (magenta, MTs), Cut11-mRFP (magenta, the NE) and LifeAct-GFP (green, actin). Corresponding kymograph images are shown on the right, in which the middle of the cell axis is shown with dotted lines and arrowheads. The numbers on the bottom right indicate times in minutes upon recording. Scale bars, 10 min (horizontal) and 5 μm (vertical). (B) Profiles of the relative position of the nucleus in *cut7-22cdc7-24* mutants in the absence (n=8) or presence (n=7) of LatA. The distance (x) between the center of the cell axis and that of the nucleus was measured at each time-point and plotted against time. (C) Classification of the patterns of nuclear movement in *cut7-22cdc7-24* cells in the absence or presence of LatA. The percentage of cells that do or do not show nuclear displacement is shown. For each sample, the maximal distance (x μm) between the center of the cell axis and that of the nucleus was determined using the data shown in (B) and categorized into three classes: displaced (shown in magenta, in which x is ≥1 μm), mildly displaced (shown in pink, in which x is between 0.5 μm and 1 μm) and centered (shown in gray, in which x is ≤0.5 μm). The data for *cut7-22* in (B) and (C) are the same as those in Figure 2B and 2C, respectively.

### *cut7* survivors become diploidized

We next asked whether the inhibition of CAR assembly/cytokinesis could rescue cut-mediated lethality of *cut7-22* cells. To this end, *cut7-22* mutants were cultured at 36°C in the absence or presence of LatA. Note that LatA not only suppresses nuclear displacement, but also inhibits CAR assembly. Remarkably, in *cut7-22* mutant cells, LatA substantially increased viability (Figure 4A). Thus, inhibition of CAR assembly and cytokinesis largely rescued lethality derived from the cut phenotype.

**Figure 4.**
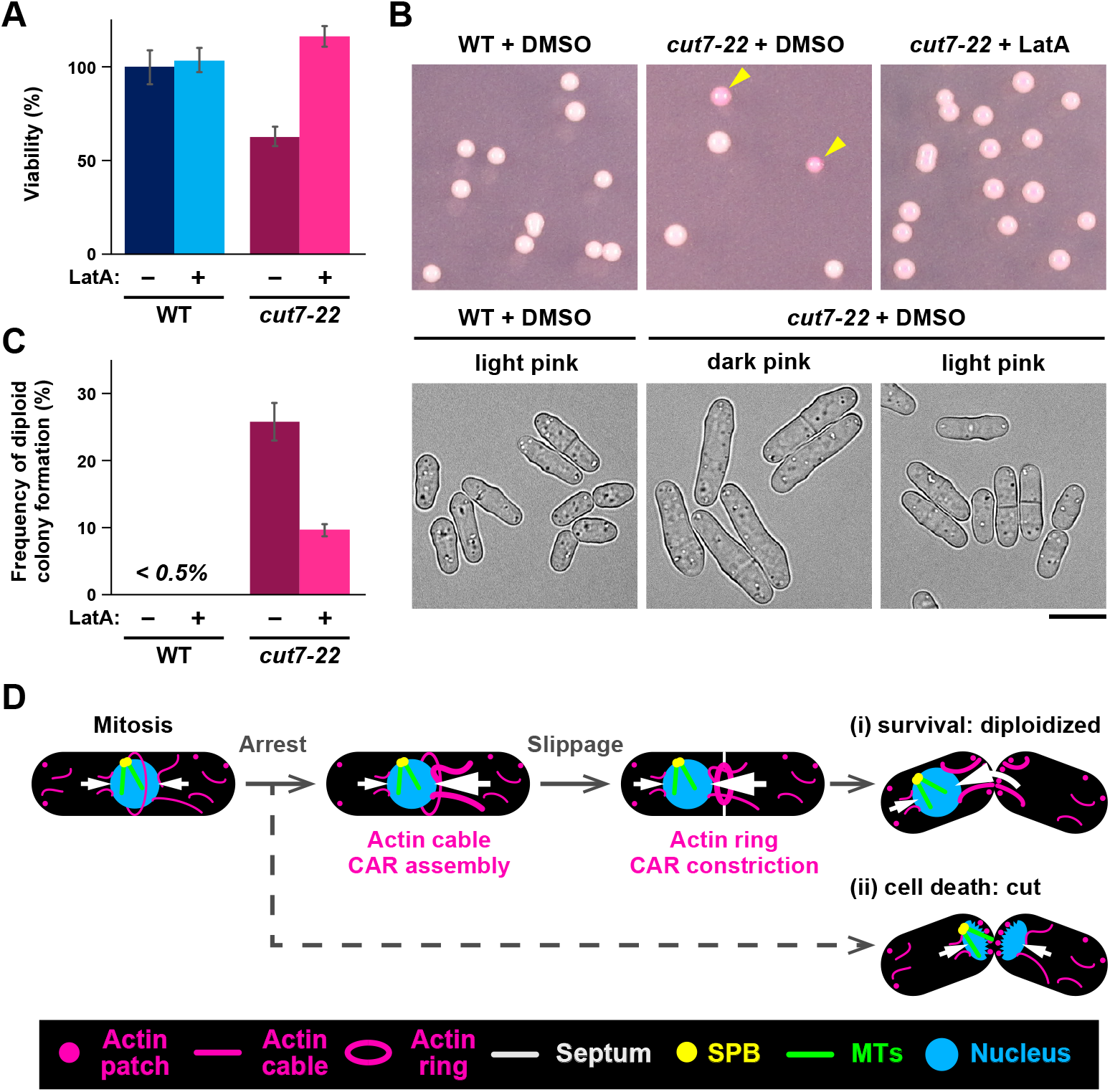
Survivors of *cut7-22* are diploidized and LatA rescues lethality of *cut7-22*. (A) Viability of wild type and *cut7-22* cells incubated at the restrictive temperature in the absence or presence of LatA. Wild type and *cut7-22* cells grown at 27°C were shifted to 36°C. After 2 h incubation at 36°C, DMSO or LatA (50 μM) was added, and cultures were incubated for additional 1 h. The cell number was measured, and after appropriate dilutions, cells were spread on rich YE5S plates containing Phloxine B. Plates were then incubated at 27°C to assess colony forming ability (viability: the number of colonies formed divided by that of cells spread, and the value of wild type cells were set as 100%). Data are presented as the means ± SEM (≥100 colonies). (B) Representative pictures of plates that contain survivor colonies (wild type, left; *cut7-22* cells, middle and right) are shown. Dark pink colonies (diploids) are pointed with arrowheads. Cell morphologies derived from light and dark pink colonies are shown on the bottom. Scale bar, 5 μm. (C) The percentage of dark pink colonies (diploid). At least 30 colonies were counted in three independent experiments. (D) A model depicting controlling mechanisms of mitotic nuclear positioning. During mitosis, actin cables nucleate in the cytoplasm and are recruited towards the cell center to be incorporated into the CAR. This process may generate a pushing force (white arrows) towards the nucleus. When mitosis is blocked, forces from either side of the nucleus becomes imbalanced, leading to nuclear displacement. Upon prolonged delay, cells are destined for two fates. In one type (i), the nucleus is further displaced from the cell center imposed by CAR constriction. Cells with displaced nuclei give rise to diploid and anucleate progenies. In another type (ii), the nucleus stays in the middle, leading to the lethal cut phenotype.

In the course of scoring the number of colonies formed at 27°C, we realized that many of *cut7-22* survivor colonies stained dark pink on Phloxine B-containing plates (Figure 4B), implying that they are diploid (Phloxine B stains diploid colonies with dark pink colors). In fact, visual inspection of cell morphology of these *cut7-22* survivors showed that these cells were wider and longer, reminiscent of diploid fission yeast cells (Figure 4B). Intriguingly, *cut7* colonies that were formed from cultures incubated in the presence of LatA were mostly haploids (Figure 4B and 4C). These results indicate that firstly the main reason for lethality of *cut7* is derived from the cut phenotype, secondly some cells could escape from cut by displacing the nucleus from the middle of the cell axis, and finally, these *cut7* survivors could resume cell division as diploid progenies at the permissive temperature. Hence, nuclear displacement provides an advantage to mitotically arrested cells for survival; however, it is brought about in exchange for genome instability.

### Conclusions

As in other organisms, actin plays a pivotal role in fission yeast mitosis; it is required for CAR assembly, a crucial step for cytokinesis. We show here that particularly upon mitotic arrest, F-actin induces the off-center nuclear positioning. Intriguingly, this nuclear displacement enables these cells to avoid lethal cut (Figure 4D). Therefore, mitotic F-actin plays a dual role; one is required for successful cell division and the other involves the protection from lethal mitotic catastrophe upon mitotic arrest and exit. It remains possible that the second role stems from a by-product of normal actin rearrangements: incorporation of centripetal mitotic actin cables into the actomyosin ring and CAR constriction.

The SAC inhibits sister chromatid separation and mitotic exit through APC/C inactivation (Musacchio, 2015). F-actin-mediated nuclear displacement appears to help evade a catastrophic consequence when the SAC is silenced and/or the septation occurred without chromosome segregation. However, this salvage is inefficient and a double-edged sword, as the nucleate compartment would resume cell division as diploids and the other anucleate part dies.

A wide range of organisms have positioned the mitotic nucleus/chromosomes to the defined locations within cells. In budding yeast mitosis, the nucleus translocates from the mother cell compartment to the bud neck, thereby positioning the mitotic spindle along the mother-daughter axis. This dynamic process requires interaction between F-actin cables and astral MTs (Caydasi et al., 2010; Fraschini et al., 2008). In vertebrate oocytes, actin diffusion centers the nucleus during prophase I and meiosis I (Almonacid et al., 2015; Almonacid et al., 2019). In human cell cultures, the actin cytoskeleton collaborates with the dynein-based machinery that exerts cortical pulling forces on astral MTs, thereby positioning the nucleus at the center of the cell (Kiyomitsu, 2019). In the moss *Physcomitrella patens*, nuclear positioning is regulated by collaborative actions between MTs and F-actin (Yi and Goshima, 2020). Whether F-actin cables directly drive movement of nucleus/chromosomes in other systems is of great interest to explore. Collectively, F-actin and its dynamics play more multifaceted roles in the maintenance of genome integrity than is currently thought.

## Abbreviations used

APC/C: Anaphase Promoting Complex/Cyclosome
CAR: Contractile Actomyosin Ring
cut: cell untimely torn
LatA: Latrunculin A
MT: microtubule
NE: nuclear membrane
SAC: Spindle Assembly Checkpoint
SPB: Spindle Pole Body
ts: temperature sensitive

## Acknowledgements

We thank Mohan Balasubramanian, Iain Hagan, J. Richard McIntosh, Paul Nurse, and NBRP (YGRC) for strains and reagents used in this study. We are grateful to Corinne Pinder for critical reading of the manuscript and useful suggestions. This work was supported by the Japan Society for the Promotion of Science (JSPS) [KAKENHI Scientific Research (A) (16H02503 to T.T.) and Scientific Research (C) (19K05813 to M.Y.)].

## Author contributions

M.Y. and T.T. designed the research. M.Y. and Y.T. performed experiments and analyzed the data. T.T. and M.Y. wrote the manuscript and Y.T. made suggestions.

## Supplemental Information

**Supplementary Table 1:**
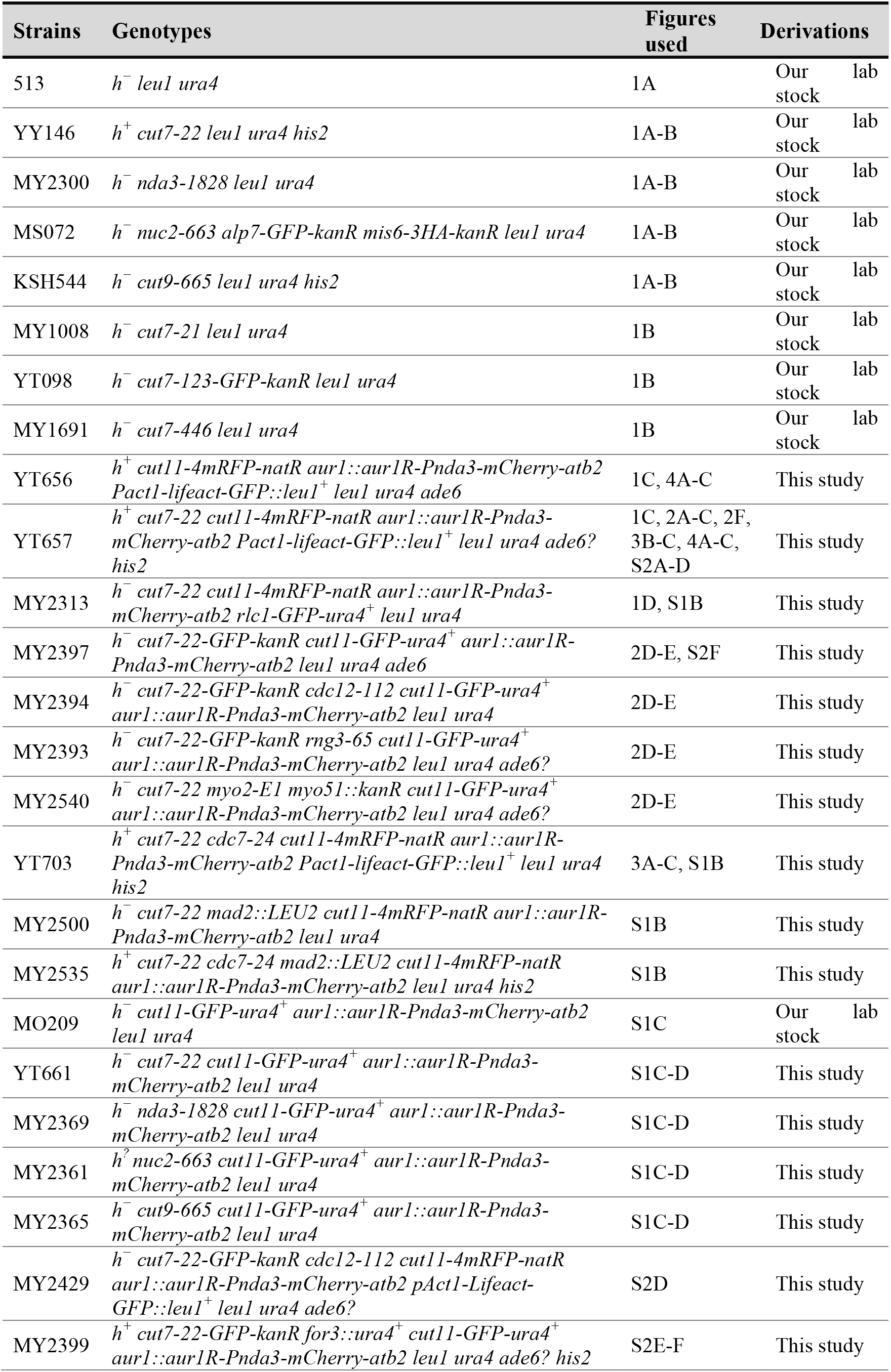
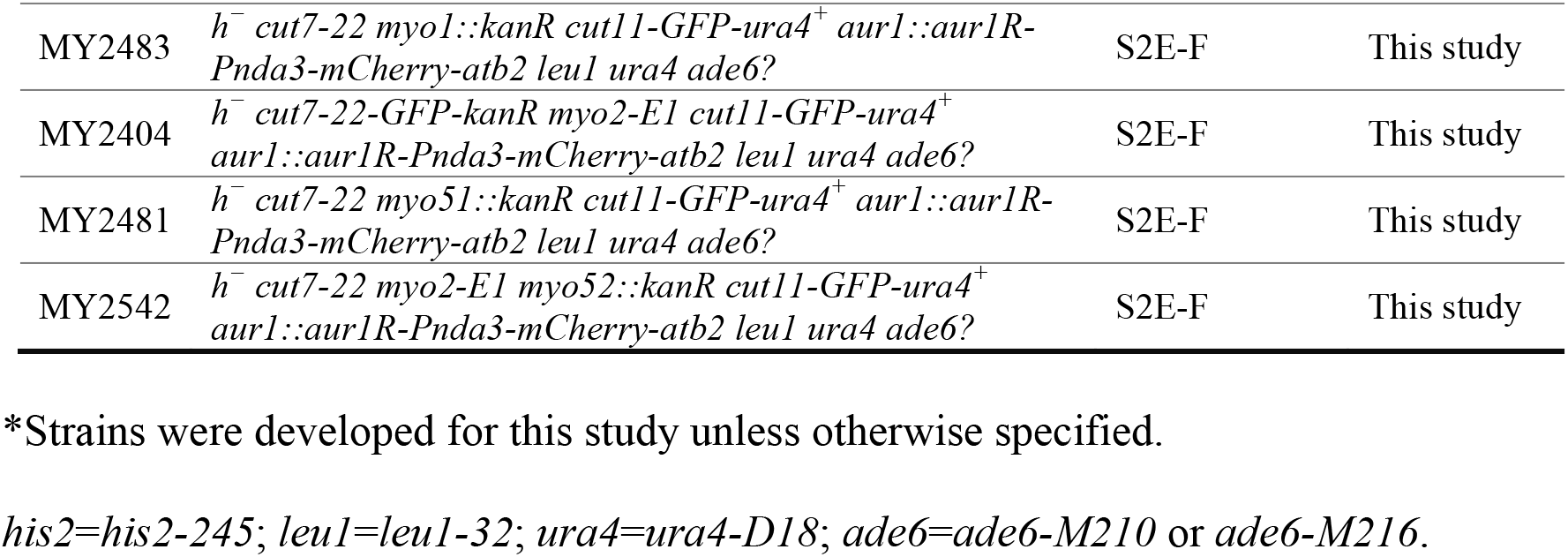
Fission yeast strains used in this study.

### Supplementary Videos

**Supplementary Video S1:** Nuclear positioning in a mitotic wild-type cell

**Supplementary Video S2:** Nuclear displacement in the *cut7-22* mutant

**Supplementary Video S3:** A *cut7-22* cell displaying cut with the nucleus being retained in the cell centre

**Supplementary Video S4:** Timing of the initiation of nuclear displacement, CAR (Rlc1) assembly and constriction in *cut7-22*

**Supplementary Video S5:** Suppression of the nuclear displacement in *cut7-22* by treatment with LatA

**Supplementary Video S6:** Suppression of nuclear displacement in *cut7-22cdc12-112*

**Supplementary Video S7:** Suppression of nuclear displacement in *cut7-22rng3-65*

**Supplementary Video S8:** Nuclear displacement in the *cut7-22 myo2-E1myo51Δ* mutant

**Supplementary Video S9:** A *cut7-22* cell in which F-actin cables interact with the nuclear membrane

**Supplementary Video S10:** Nuclear displacement in *cut7-22cdc7-24*

**Supplementary Video S11:** Suppression of the nuclear displacement in *cut7-22cdc7-24* by treatment with LatA

### Supplementary Figure Legends

**Figure S1.**
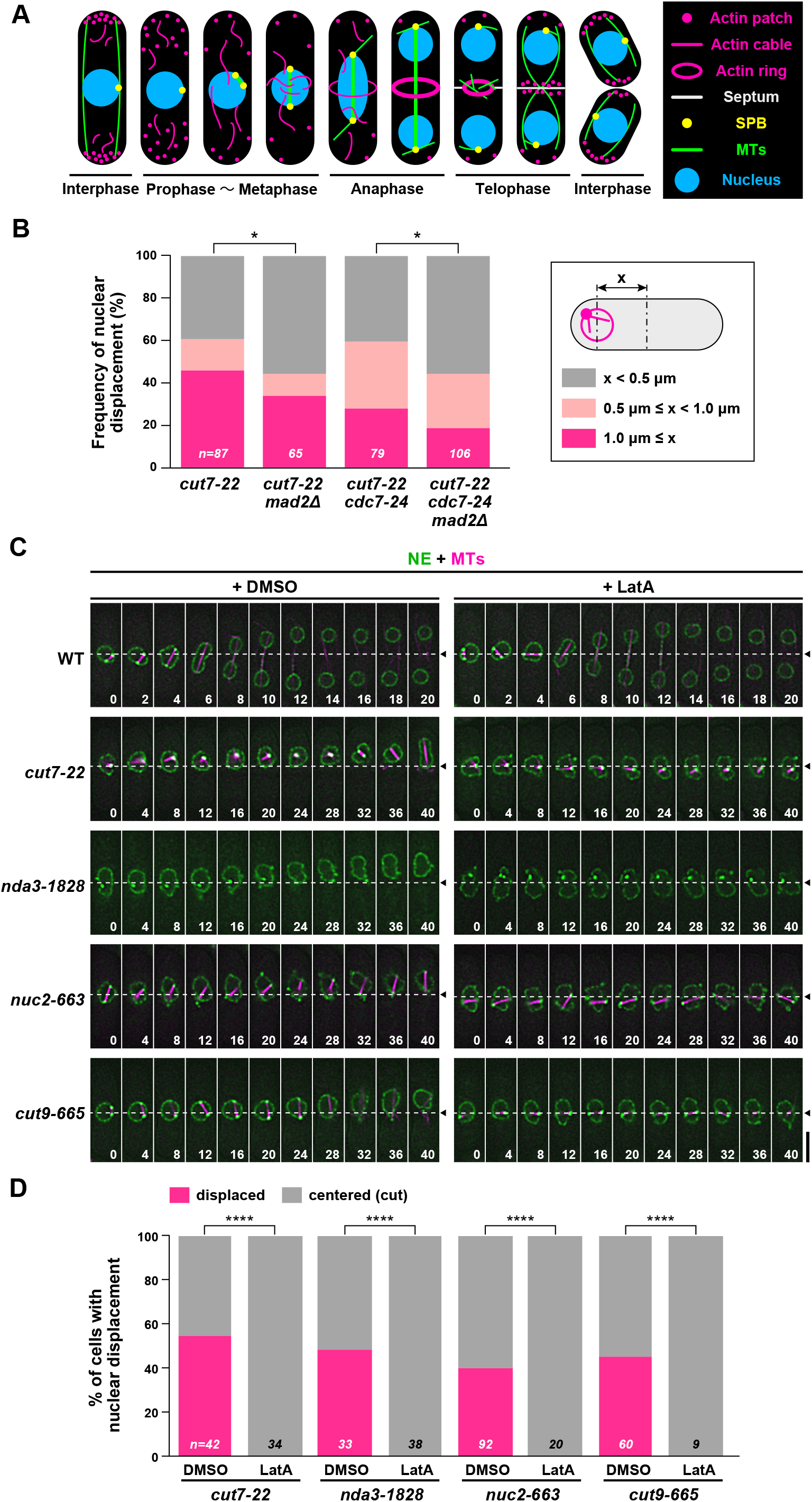
Impact of the spindle assembly checkpoint and suppression of nuclear movement with depolymerization of the F-actin cytoskeleton. Related to Figure 1. (A) A schematic depiction of the nucleus and cytoskeletons during the mitotic cell cycle in fission yeast. (B) The degree of nuclear movement in *cut7-22* or *cut7cdc7-124* mutants in the presence or absence of *mad2*. The percentage of cells that do or do not show nuclear displacement is shown. For each sample, the maximal distance (x μm) between the center of the cell axis and that of the nucleus was determined and categorized into three classes: displaced (shown in magenta, in which x is ≥1 μm), mildly displaced (shown in pink, in which x is between 0.5 μm and 1 μm) and centered (shown in gray, in which x is ≤0.5 μm). The sample number is shown on the bottom of each column. (C) Time-lapse images of the nuclear position in indicated mutant cells in the absence or presence of LatA. Wild-type and individual mutant cells grown at 27°C were shifted to 36°C and incubated for 2 h except for *nda3-1828*, which was incubated for 5 h. DMSO or LatA (50 μM) was then added and cells were incubated for further 10 min, when imaging started (time 0). Numbers on the bottom right corners of each image show times in minutes. Cells contained mCherry-Atb2 (magenta, MTs) and Cut11-GFP (green, the NE). The middle of the cell axis is shown with dotted lines and arrowheads. Scale bar, 5 μm. (D) Profiles of nuclear positioning in individual mitotic mutants in the absence or presence of LatA. For each strain, the maximal distance (x μm) between the center of the cell axis and that of the nucleus at each time point was determined. If x was kept <1 μm for longer than 20 min or a cell showed cut, it was classified as centered (cut). If x was >1 μm during observation, it was classified as displaced. The sample numbers (n) are shown on the bottom for each column. All *p*-values were obtained from the two-tailed χ^2^ test: * *p* <0.05; **** *p* <0.0001.

**Figure S2.**
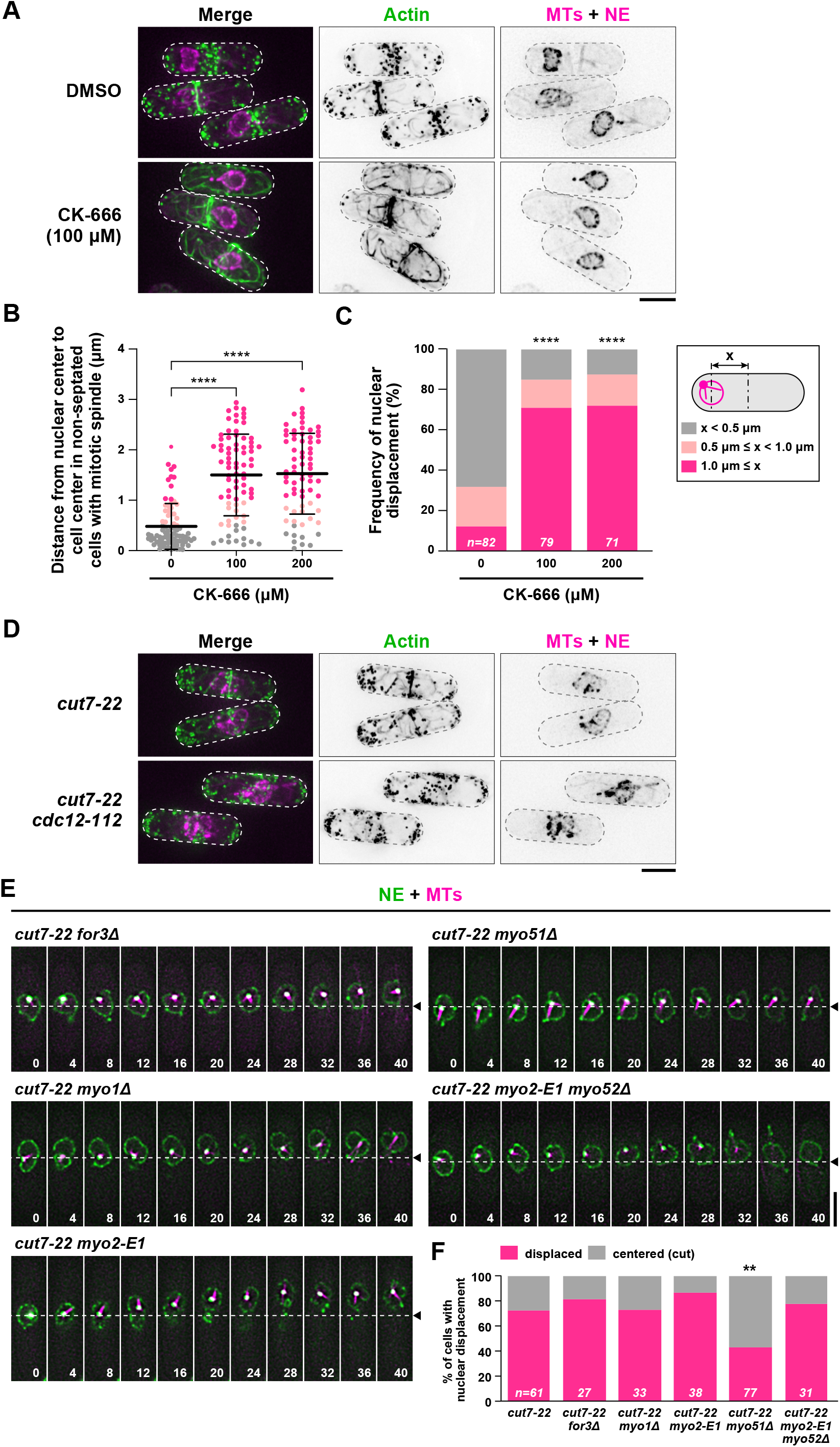
Enhanced nuclear movement in the presence of CK-666 and the involvement of Myo51. Related to Figure 2. (A) Lack of actin patches in *cdc7-22* cells treated with CK-666. *cut7-22* cells containing mCherry-Atb2 (MTs, magenta), Cut11-mRFP (the NE, green) and LifeAct-GFP (actin, green) were grown at 27°C and shifted to 36°C for 2 h. CK-666 (100 μM) was added and images were taken 20 min after CK-666 addition. Scale bar, 5 μm. (B) The nuclear position in *cut7-22* cells treated with CK-666. *cut7-22* cells containing mCherry-Atb2 (MTs), Cut11-mRFP (the NE) and LifeAct-GFP (actin) were grown at 27°C and shifted to 36°C for 2 h. CK-666 (100 or 200 μM) or DMSO was added. Samples were observed 20 min after CK-666 addition. The distance (x μm) between the center of the nucleus and that of the cell center was measured for each cell and plotted in the graph as dots: gray, x<0.5 μm; pink, 0.5 μm< x<1.0 μm; magenta, x>1.0 μm. Mitotic cells prior to CAR constriction were counted for nuclear positioning. Data are given as mean±s.d. (C) Patterns of nuclear position. Data shown in (A) are presented as columns. (D) Lack of actin cables and the CAR in *cdc7-22cdc12-112* cells. *cut7-22* or *cut7-22cdc12-112* cells containing mCherry-Atb2 (MTs, magenta), Cut11-mRFP (the NE, green) and LifeAct-GFP (actin, green) were grown at 27°C and shifted to 36°C for 4 h. Scale bar, 5 μm. (E) Time-lapse images of the nuclear position. Indicated mutant cells grown at 27°C were incubated at 36°C for 2 h, when live imaging started. Individual mutants contain mCherry-Atb2 (MTs, magenta) and Cut11-GFP (NE, green). The position of the cell center is shown with dotted lines and arrowheads. Scale bar, 5 μm. (F) Profiles of the relative position of the nucleus. If x (as in (C)) was <1 μm or a cell showed cut, it was classified as centered (cut). If x was >1 μm, it was classified as displaced (magenta). The sample numbers are shown on the bottom of each column. All *p*-values were obtained from the two-tailed χ^2^ test: ** *p* <0.01; **** *p* < 0.0001.

